# Machine learning-based detection of insertions and deletions in the human genome

**DOI:** 10.1101/628222

**Authors:** Charles Curnin, Rachel L. Goldfeder, Shruti Marwaha, Devon Bonner, Daryl Waggott, Undiagnosed Diseases Network, Matthew T. Wheeler, Euan A. Ashley

## Abstract

Insertions and deletions (indels) make a critical contribution to human genetic variation. While indel calling has improved significantly, it lags dramatically in performance relative to single-nucleotide variant calling, something of particular concern for clinical genomics where larger scale disruption of the open reading frame can commonly cause disease. Here, we present a machine learning-based approach to the detection of indel breakpoints called *Scotch*. This novel approach improves sensitivity to larger variants dramatically by leveraging sequencing metrics and signatures of poor read alignment. We also introduce a meta-analytic indel caller, called *Metal,* that performs a “smart intersection” of Scotch and currently available tools to be maximally sensitive to large variants. We use new benchmark datasets and Sanger sequencing to compare *Scotch* and *Metal* to current gold standard indel callers, achieving unprecedented levels of precision and recall. We demonstrate the impact of these improvements by applying this tool to a cohort of patients with undiagnosed disease, generating plausible novel candidates in 21 out of 26 undiagnosed cases. We highlight the diagnosis of one patient with a 498-bp deletion in *HNRNPA1* missed by traditional indel-detection tools.

## Background

Insertions and deletions within the genome are well-established mechanisms of human disease^1^. While less common than single-nucleotide variants, indels are an important component of genetic diversity, and are more likely to disrupt the open reading frame^2,3^. In a recent version of the Human Genome Mutation Database (HGMD 2018.4) indel mutations account for 31% of all entries, with deletions outnumbering insertions more than 2:1^4^.

While single-nucleotide variants (SNVs) can be accurately detected using next-generation DNA sequencing and relevant software, indel detection remains a challenge^5^. Previous studies found low concordance between indel calls made by commonly used indel-calling pipelines or sequencing platforms^6,7^.

By comparison with long-read sequencing, indel calling from short-read sequencing has been shown to miss variants, including clinically relevant ones^8^. The current diagnosis rate through exome sequencing for patients with genetic disorders is around 30%^9,10^; one possible explanation for this low diagnosis rate is that the genetic cause of many patients’ diseases are a type of variant that are difficult to detect, including indels.

Since indels are a crucial source of genetic diversity that are known to cause disease, it is important to understand how currently available indel detection tools perform. Evaluating the performance an indel caller involves comparing the variants it identifies against a “benchmark” or “truth” set of variants that we accept as fully characterizing the variation of the genome (possibly within certain genomic regions and/or categories of variation). This process produces scores of recall (sensitivity) and precision (positive predictive value), derived from the proportions of true positive, false negative, and false positive calls. These metrics capture how likely a genuine variant is to be called, and how likely a called variant is to be genuine. The ideal indel caller has high recall and high precision, reporting many genuine variants and few false ones; it moreover detects a variety of indels, including variants of different sizes, and those that lie in different types of genomic regions (e.g., homopolymer runs).

Given a truth set for a benchmark genome and the query set of variants reported by an indel caller in the same, we perform a breakpoint-based comparison to generate metrics describing the caller’s performance. We subdivide each list into sublists of insertion breakpoints, deletion start breakpoints, deletion end breakpoints, and breakpoints of any type. Corresponding truth-query pairs are input to an app developed by the Global Alliance for Genomics and Health (GA4GH Benchmarking) made available on precisionFDA^11^. The benchmarking tool then classifies calls as true positives, false positives, or false negatives. True positives are variants in the truth set that are within 3 bp of a call in the query. False negatives are the remaining calls in the truth set that are not within 3 bp of such a call. False positives are variants in the query set that are not within 3 bp of a call in the truth set. The proportions of calls in these categories are used to to derive precision and recall scores.

While not explicitly a part of the comparison, this method does credit or penalize tools for their estimate of the size of a deletion, which determines where the caller places its start and end breakpoints. It does not rely on zygosity or alternate allele sequences, which Scotch and Pindel do not report for all variants.

This process produces recall and precision scores for each class of breakpoint. For deletions, we report the mean metric across start and end breakpoints. For each tool, we also benchmarked the full set of all called indel breakpoints against all true positive breakpoints. This illustrates a caller’s performance on both insertions and deletions, with an emphasis on performance on deletions, since in sequencing data each deletion comprises two breakpoints (start, end) while an insertion has only one. Combining all breakpoints also expresses a tool’s ability to determine that an indel of some sort exists at a given locus, even if the type is not correctly identified.

We separate recall into two metrics. Recall by count is the proportion of the number of true positive breakpoint calls, to true positive and false negative breakpoint calls. Recall by base is the proportion of bases belonging to indels with true positive breakpoint calls, to bases belonging to indels with true positive and false negative calls. The latter highlights a caller’s sensitivity to larger variants. GA4GH Benchmarking reports recall by count directly; we calculate recall by base by considering the number of bases belonging to true positive and false negative variant calls.

Benchmark data derived from finding consensus between multiple orthogonal variant callers is often labeled “gold standard.” These variant callers may not individually detect all true positives in the genome; moreover, while the requirement of including only variants identified by multiple callers ensures that the variants in the benchmark set are of high confidence, it may leave out additional genuine variants. Incompleteness of benchmark datasets can warp benchmarking metrics, flagging real calls made by callers outside the truth set as false positives (deflating estimates of precision). The decreased number of true positives may also lead to inflated estimates of recall. Additionally, we note that machine-learning approaches trained on an incomplete benchmark dataset are limited, learning to classify real indels missing from the truth set as normal loci.

There are currently several categories of indel callers that detect different types of indel signatures. Split-read tools, such as SV-M^12^, extract read pairs where only one mate can be confidently mapped to the reference, and align the unmapped mates. Paired-end tools, like BreakDancer^13^, examine the distribution of insert sizes between mate reads in a given region to determine whether sequence has been added or removed. Alignment-based tools like Stampy^14^ examine how reads align to a reference. Some alignment-based tools, for example Platypus^15^, may conjecture alternate haplotypes to which reads are aligned. Others, such as Scalpel^16^, perform graph-based alignments. For sensitive detection of larger indels, there are fewer choices. The most popular tool may be Pindel^17^, which uses a pattern-growth algorithm to detect indels through split reads. Other tools, like IMSindel^18^ and ScanIndel^19^, use *de novo* assembly to identify large variants.

Smaller indels (1 to 5 bp) make up a majority (77%) of indels by number, but they account for a very small proportion (6%) of all inserted and deleted bases in the human genome. By number of nucleotides, larger indels exert significantly more influence on the diversity of the genome. But many callers struggle to identify larger indels. Applied to simulated variants of up to 50 bp in length, popular callers such as GATK UnifiedGenotyper^20^, SAMTools^21^, and VarScan^22^ detected no indels greater than 37, 44, and 42 bp in length, respectively^23^.

A distinct category of tools exists for detecting copy-number variants (CNVs) and structural variant (SVs), large-scale genetic abnormalities of a kilobase or more in length. But this leaves few options for sensitive detection of indels larger than a few bases and smaller than one kilobase. Insertions, which cannot be detected through changes in sequencing depth, can be particularly challenging. Our goal was to build on the capabilities of recent tools and leverage the availability of improved benchmark datasets to develop an indel caller with increased sensitivity to variants across the size spectrum.

## Results

### A benchmark genome of simulated indels facilitates evaluation

One source of benchmark data is simulated variants. The primary advantage presented by this technique is that the truth set is known with maximal confidence, since variants are precisely “spiked in.” This improves the reliability of calling metrics. Additionally, the incidence of different kinds of indels can be manipulated to generate sufficient test data to characterize indel callers’ performance on a variety of variants. The primary drawback of simulated indels is that they may not perfectly recapitulate real genotypes.

There exist several tools that can be precisely directed to spike variants into a starting genome. Using BAMSurgeon^24^, we defined 3,885 non-overlapping indels, three with each possible size from 5 to 1000 bp—and then one indel for each 10 bp increment between 1000 and 10000 bp, on average. Half were deletions, and half were insertions for which the inserted sequence was generated randomly. The positions for these variants were selected randomly from mappable regions of chromosome 22 at least 10 bp away from known pre-existing indels in a starting genome (NA12878). When benchmarking variant callers’ performance on this dataset, we also excluded the known pre-existing true positives in NA12878 and calls made by callers within 10 bp of them. We note the benchmark-incompleteness issue still exists here: variants in NA12878 but missing from its truth set will be present in this dataset, and callers’ capture of them may incorrectly be flagged as false positives, deflating estimates of precision.

A minority of these variants (n = 860) were rejected by BAMSurgeon because they could not be made to appear biologically realistic, usually because starting coverage was too low, or a sufficiently large contig into which to inject the variants could not be assembled. (Above 8,000 bp in length, we found, it was increasingly unlikely that the indels could be introduced.) We first used this dataset—which is distinct from the simulated variants used to supplement the training data of our new tool, though made by the same simulation software—to benchmark the variant callers.

### Syndip offers a comprehensive set of variants for benchmarking

“Gold standard” datasets available for benchmarking include public genomes such as NA12878^25^ and Syndip^26^. NA12878 is the genome of a woman from Utah, whose variants are characterized by the Genome in a Bottle Consortium (GiaB) according to consensus calling across multiple sequencing and variant calling platforms. Syndip is a recently released synthetic diploid genome produced by combining two haploid human cell lines sequenced using single molecule real-time sequencing and identifying indels with FermiKit^27^, FreeBayes^28^, Platypus^15^, Samtools^29^, GATK HaplotypeCaller and GATK UnifiedGenotyper^30^ (Supp. Fig. 4).

Relative to NA12878, Syndip offers a wider range of indels that provide for more comprehensive benchmarking. Syndip includes more indels than NA12878, and these variants span a greater range in size. While across NA12878, the mean and standard deviation of indel sizes is just 3 and 4 bp, in Syndip it is 22 and 209 bp. And while in NA12878, the largest variant is just 127 bp, in Syndip, it is 19 kb. Syndip was also developed with long-read sequencing, which incurs random errors that can generally be overcome by sequencing depth. Neither of the two machine learning-based callers evaluated here (DeepVariant and Scotch) were trained on Syndip. GATK HaplotypeCaller may have an advantage as it was one of the original tools used to develop the Syndip truth set.

### A machine-learning based caller designed for capturing large indels

We present a machine learning-based tool focused on increasing the sensitivity of calling larger indels from whole-genome sequencing data (Scotch, Fig. 1). Machine-learning techniques, including random-forest modeling^31^, have been applied to variant calling with success before. DeepVariant^32^, which uses neural networks to analyze pileups, won highest “SNP Performance” in the precisionFDA Truth Challenge. Scotch examines designated portions of the genome, and analyzes each base individually. It creates a numerical profile of these positions, describing various features of the aligned sequencing data like depth, base quality, and alignment to the reference. A full explanation of the features selected is available in the Supplementary Note.

**Fig. 1:**
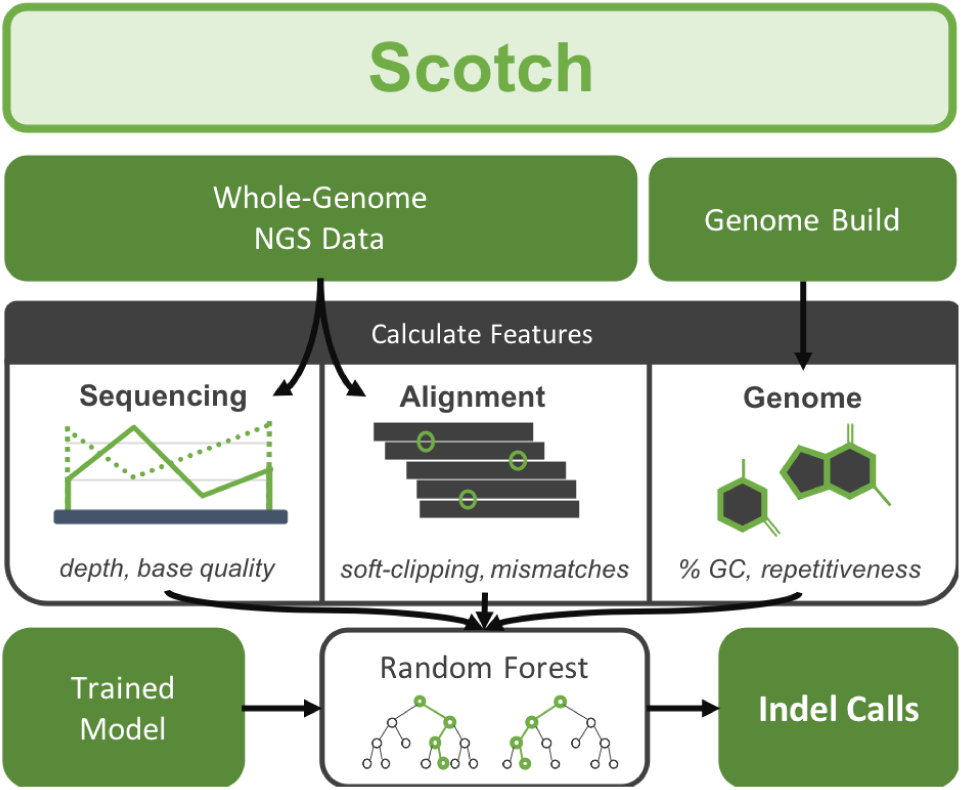
Scotch is a machine-learning based indel caller. Features are calculated from input sequencing data and from a reference genome. A random forest identifies positions that are the breakpoints of an insertion or deletion.

Based on these predictors, a random forest model then classifies the position as non-indel or the site of a specific type of indel breakpoint. If the locus does not match the reference, Scotch classifies it as one of three types of indel breakpoints—the site of an insertion, the start of a deletion, or the end of a deletion—or as a 1-bp deletion, which requires a separate class since both deletion breakpoints fall on the same locus, considered in half-open notation. The Scotch standard model was trained on the NA12878 genome. We added to its training data larger simulated indels as a way of attempting to overcome the incompleteness of the benchmark dataset, which generally lacks larger indels.

We evaluated Scotch and five other callers: DeepVariant; GATK HaplotypeCaller^30^; VarScan2^22^; and two versions of Pindel, the standard, which we refer to as “Pindel”, and the pipeline with the “-l” option for reporting long insertions, “Pindel-L”. These versions of Pindel call the same deletions; Pindel-L includes many additional insertions. We assess the performance of these six pipelines on three benchmark datasets: simulated variants, Syndip^26^, and NA12878^25^. For ease of use, we subset chromosome 22 from each of these datasets. This includes approximately 3000, 9900, and 8700 indel breakpoints, respectively, with each deletion contributing two breakpoints and each insertion contributing one. The full results of this benchmarking are available in Supplementary Tables 1 - 9. Below, we concentrate primarily on the simulated variant dataset that contains many large variants, and Syndip, which offers the most comprehensive set of variants.

### Scotch has high recall on simulated data, identifying indels of up to several kilobases

Evaluated on the dataset of simulated indels, most callers perform well in identifying small variants. However, their performance generally declines markedly as indel size increases (Fig. 2). In contrast, across all indel breakpoints, Scotch’s recall by count and recall by base (99%; 99%) both exceed Pindel-L (79%; 74%), which itself surpasses all other tools. On deletions, Scotch (recall by count: 97.9%), is incrementally superior to Pindel (97.2%), and far exceeds other individual callers (which range in recall from 1% to 12%). On insertions, the differences are even more clear: Scotch (recall by count: 98%) surpasses Pindel-L (42%) and all other individual tools (0.3% - 24%). Scotch retains high recall across the size spectrum, successfully identifying insertions and deletions in the dataset larger than the largest variant on which it was trained (500 bp), and including the dataset’s largest insertion of 7810 bp and largest deletion of 7608 bp.

**Fig. 2:**
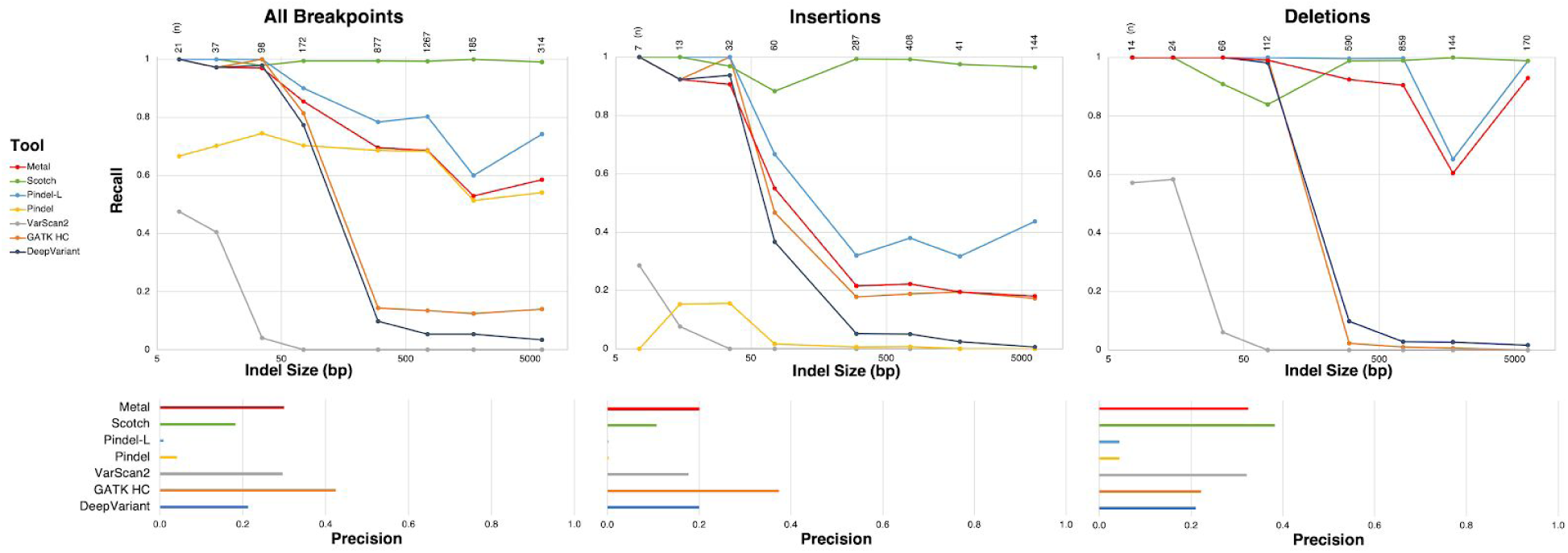
Recall by indel size on simulated variants. To examine how callers’ performance varies with indel size, we group together variants with sizes within a fixed range and assess the caller’s sensitivity to the group. The performance of many popular callers declines as indel size increases. Scotch’s performance, in contrast, is more consistent across the size spectrum. (Note: Since Pindel and Pindel-L call the same deletions, their recall curves on these variants are overlapping.) The simulated dataset, produced by excluding known indels in the NA12878 genome and spiking in a smaller number of indels into the same genome, produces deflated estimates of precision. (It includes all the loci at which a tool would make a false positive call in NA12878, while offering a smaller number of true positive variants.) This data is also available in Supplementary Tables 16 - 18.

Across this broad range of indel sizes, we note that the full relationship between a caller’s recall and indel size is complex. While, in general, as indel size increases, sensitivity decreases, there are important exceptions. Pindel-L registers a sharp decline, then increase in recall on deletions around 1 kb. For most callers, the steepest drop in recall occurs near 150 bp, approximately the length of a short read. These variants may be particularly difficult to detect, because unlike smaller variants, they cannot be contained in a single short read. Beyond 150 bp, recall is still somewhat variable, and some callers continue to decline in performance while others rebound slightly.

### Scotch identifies variants in Syndip with high recall

Similar performance is seen when these tools are tested on Syndip (Supplementary Tables 1 - 3). Across all indel breakpoints, Scotch’s recall by count is the highest of any tool (93%), exceeding Pindel-L (91%), DeepVariant (87%), GATK HaplotypeCaller (87%), VarScan2 (73%), and Pindel (66%). Scotch and Pindel-L, furthermore, call variants that account for 68% and 71% of all inserted and deleted sequence, respectively. In contrast, the variants identified by DeepVariant, GATK HaplotypeCaller, and VarScan2 account for between 15% and 36% of all inserted and deleted sequence.

Scotch’s high recall is attributable to the features included in the random forest model. While Scotch does leverage some features from alignment that other tools use too (e.g. aligning with an insertion or deletion against the reference), Scotch also utilizes features that many other tools ignore, such as the proportion of reads at a given location that have been soft clipped. More information on the features is available in the methods section, with a full list of features and their importance (as defined by the mean decrease in Gini Index) available in Supplemental Figure 2 and Supplemental Table 19.

### Scotch has low precision on consensus benchmark data

While Scotch has high recall, on consensus data sets, it exhibits low precision. By precision, Scotch (35%), Pindel (27%) and Pindel-L (6%), fall far behind VarScan2 (98%), DeepVariant (95%), and GATK HaplotypeCaller (91%). Estimates of Scotch’s precision, however, may be deflated by its identification of real variants missing from the truth set. As discussed below, Sanger sequencing of Scotch’s calls flagged in benchmarking as false positives reveals that many are *bona fide* variants. Note also the differential metrics: Scotch’s insertion-specific precision is 21%, while its deletion-specific precision is 76%.

### Metal: a meta-analytic indel caller sensitive to large variants

Each of the benchmarked callers has its own strengths, and none outperforms all others in all circumstances. To produce an optimal variant calling tool, we merge their strengths. Integrating multiple variant callers into a meta caller has been shown to improve performance^33,34^.

Some tools achieve excellent performance in one dimension by sacrificing performance in another. VarScan2, for example, is very conservative, and thus attains extremely high precision by accepting low recall. Here, instead, we attempt to negotiate a “best compromise.” This approach, which we call Metal, retains the high precision of DeepVariant, GATK HaplotypeCaller, and VarScan2, while incorporating many of the larger variants that Scotch and Pindel-L identify. It achieves this by performing a “smart intersection.” Metal will report a call produced by a tool if it has a corresponding call within 3 bp identified by another tool. Metal is available on GitHub at https://github.com/AshleyLab/metal. To counter the low insertion-specific precision of Scotch and Pindel-L, we require that insertions called by these tools have correlates in higher-precision DeepVariant, GATK HaplotypeCaller, or VarScan2. Metal does not consider the various quality scores that tools report with the variants they call, which may not be directly comparable because tools use different scales, but integrating this evidence is a promising area for further improvement. An additional machine learning model, in fact, could arbitrate the calls made by each tool and collate them into a single set with maximal confidence.

Across all indel breakpoints in Syndip, Metal’s recall by count (90.2%) surpasses all tools except Pindel-L (90.6%) and Scotch (93.4%), while providing a 15x increase in precision relative to Pindel-L (Metal: 89%; Pindel-L: 6%), and a 2.5x increase in precision relative to Scotch (Scotch: 35%). Metal identifies far more large variants than DeepVariant, GATK HaplotypeCaller, or VarScan2, with a recall by base of 56%. On deletions, the performance benefits are particularly clear. Metal identifies more variants by count than any individual tool (90%), and more by base (60%) than all tools except Pindel and Pindel-L, with greatly improved precision (84%) compared to Scotch and versions of Pindel.

Judged by F1 score on Syndip, Metal surpasses all traditional callers; its F1 with recall by count (89%) is somewhat behind DeepVariant (91%), while its F1 with recall by base (69%) far exceeds all other tools (next best, DeepVariant: 52%). And with increasing priority given to recall, the impact of Scotch and Metal’s superior recall becomes clear (Fig. 4).

**Fig. 3:**
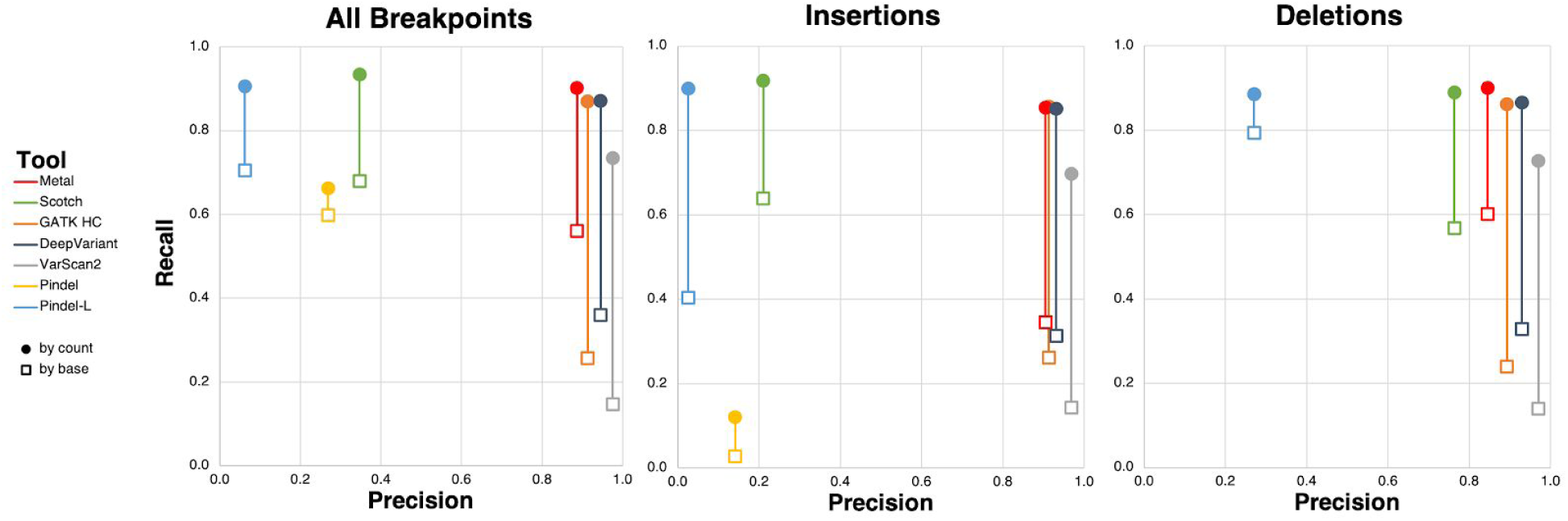
Scotch and Metal offer high sensitivity to large variants and improved precision in Syndip. Performance of the selected indel callers, including Scotch and the meta caller Metal, on Syndip. Recent callers such as DeepVariant and GATK HaplotypeCaller have married high precision and high recall by count, but they are more likely to miss larger variants. Scotch and Pindel-L offer higher recall, especially on a per-base basis, but with lower precision. Metal, a combination of the other pipelines captures some of the breakpoints of large variants Pindel-L and Scotch detect while sacrificing little in precision.

**Fig. 4:**
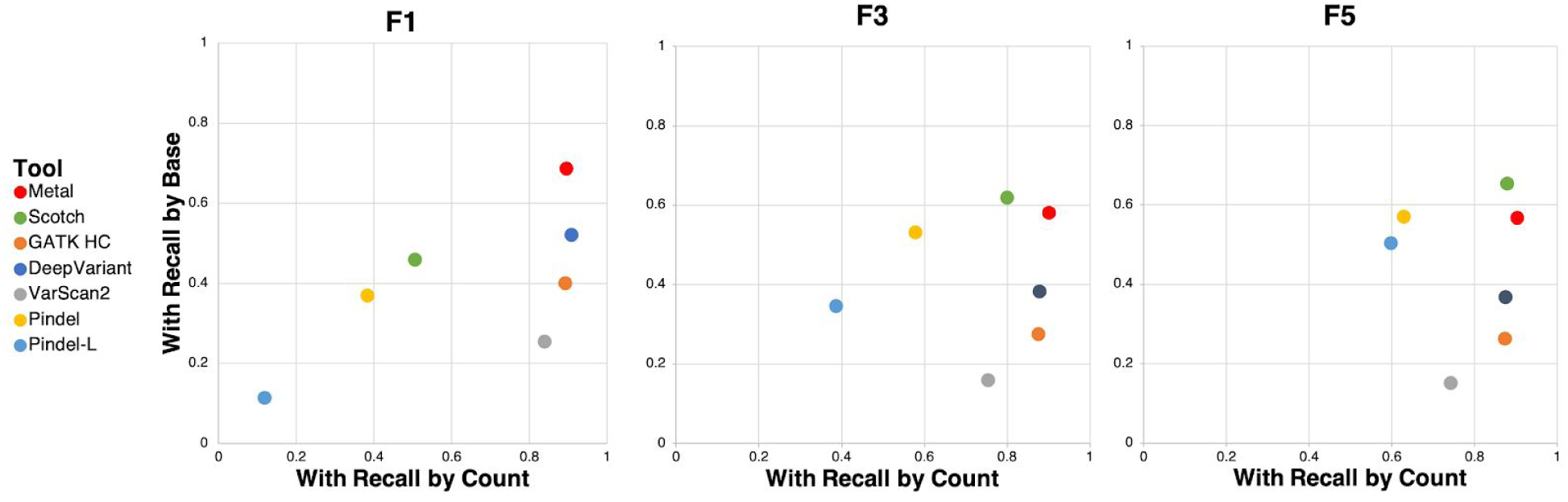
F scores with recall by count and recall by base in Syndip. We plot the F scores of the tools examined. In an F(N) score, recall is considered N times more important than precision. Recent callers such as DeepVariant and GATK HaplotypeCaller provide higher precision but are more likely to miss large variants. As the weight given to recall grows, Scotch and Metal surpass other callers.

### Sanger sequencing validates variants not in consensus truth set, altering precision estimates

While more successful than other tools in identifying larger variants, Pindel-L, and, to a lesser extent, Scotch, exhibit low precision on the consensus truth set. On Syndip, the precision of DeepVariant, GATK HaplotypeCaller, and VarScan2 lies between 89% and 97% for deletions and 91% and 97% for insertions. On deletions, Scotch’s precision is lower (76%), but higher than Pindel-L (27%) while on insertions, both Scotch (21%) and Pindel-L (3%) decline significantly.

We carried out Sanger sequencing to determine whether variants identified as false positives by Scotch were, in fact, real variants absent from the consensus truth set (see Methods for more details on how these were selected).

We Sanger selected for sequencing 100 breakpoints called by Scotch and flagged as false positives, with two constraints. First, half of the selected calls were insertion breakpoints, and half were deletion breakpoints (which, in turn, were half deletion-start and half deletion-end breakpoints). Second, 20 of the 100 calls were selected to have potential correlates in Syndip: indel calls within 3 bp, of any type, in the Syndip truth set. This constraint was introduced to determine whether Syndip had identified any additional common indels not detected by NA12878.

We analyzed the resulting chromatograms with Poly Peak Parser^35^, an online alignment-based tool that identifies indel. (The full results are available in Supplementary Table 20.) For 26 of the original 100 calls, surrounding GC content had been too high for effective primer design or PCR amplification failed. In an additional 20 cases, sequencing quality was too low to make an accurate determination. We further excluded 2 calls that were flagged as false positives not because they were missing altogether from the NA12878 truth set, but because Scotch had mis-identified their type. 18 out of the remaining 52 calls were verified as genuine indel breakpoints. 14 are homopolymer deletions.

This indicates that some variants identified by Scotch flagged as false positives when benchmarking are genuine indels. Of the 25 deletions sequenced, 12 were validated as real deletions. Of the 27 insertions sequenced, 6 were validated as real variants — 5 as deletions, and 1 as an insertion. Scotch had called 2107 deletion breakpoints and 14895 insertion breakpoints in chromosome 22 of NA12878 that were flagged as false positives.

The Sanger validation rates entail some uncertainty because of the small size, and these samples may not be perfectly representative of the full population of putative false positive results due to the constraints defined in selection. But they indicate that hundreds of Scotch’s reportedly false positive calls do indeed refer to real variants missing from chromosome 22 of the NA12878 GiaB truth set. This result indicates that estimates of Scotch’s precision are deflated.

### Scotch identifies clinically relevant variants missed by other tools

Our approach to improving indel calling was motivated by clinical application. While exome and genome sequencing are effective in diagnosing rare genetic disorders, estimates of diagnosis rate using this technology fall between 30% and 40%, leaving many patients undiagnosed. With Scotch, we sought to develop a method that would find these presumed genetic causes identifying as many true positives as possible, while minimizing the risk of missing the variant of interest.

We applied Scotch to the genomes of several patients presenting to Stanford’s Undiagnosed Diseases Network (UDN). The UDN is a national consortium of medical centers taking on patients whose atypical constellations of symptoms have evaded diagnosis. We chose a representative sample of undiagnosed patients at the Stanford center and applied the Scotch algorithm. (General information about these cases, including age, sex and phenotype terms, is available in Supplementary Table 21.) In 21 of 26 cases, plausible candidates for the diagnosis were found. Such candidates would require further work to firmly establish disease causality but their presence in a large majority of undiagnosed cases is encouraging, especially considering that most variants detected in this way could be assumed to contribute to at least hemizygous loss of function. One example case illustrates the power of the new approach well.

An adult woman presented with distal asymmetric myopathy including scapular winging, mild facial weakness, decreased forced expiratory volume, and muscle biopsy notable for rimmed vacuoles and myofibrillar disorganization. In addition to a myopathy gene panel that was negative, whole-exome sequencing was performed for the patient, without a diagnosis. With whole genome sequencing data, Scotch made 4.5m indel breakpoint calls. (This is more than VarScan2, GATK HaplotypeCaller, and DeepVariant (1.1 - 1.9m), but fewer than Pindel (5.2m) and Pindel-L (12.7m).) Of Scotch’s calls, 4,365 were deletion breakpoints within 100 bp of exons of ClinVar- and OMIM-annotated genes. 460 of these were seen in no unrelated samples, and a phenotype-based prioritization tool^36^ ranked breakpoints of a 498-bp exon-skipping stop-loss deletion in *HNRNPA1* in rank 50, which was orthogonally confirmed by quantitative PCR. This deletion was not reported by DeepVariant, GATK HaplotypeCaller, or VarScan2; it was identified by Pindel, Pindel-L, and Metal. The implicated gene is a member of the hnRNP family, which has important roles in nucleic acid metabolism; mutations in *HNRNPA1* have been previously implicated in neuromuscular disease in patients with features which substantially overlap our case’s phenotype.

## Discussion

Here, we present an approach for improved detection of insertions and deletions, called Scotch. This has applications for understanding genetic diversity and improving the diagnostic rate, using genetic testing, in the clinic. Note that Scotch prioritizes recall over precision. While low precision is addressed in clinical pipelines through filtering steps and manual curation that eliminate out false positives, low recall means losing variants that cannot be recovered.

Evaluating Scotch on the Syndip dataset, we found that Scotch has higher sensitivity than any other indel detection tool benchmarked here. Scotch reports variants previously only accessible to long-read sequencing. Evaluated on simulated data, Scotch retains recall by count close to 1 on variants across the size spectrum. The meta-calling approach incorporating Scotch surpasses all individual tools overall and is likely to improve the diagnostic rate of clinical genome calling.

Note that indel detection is highly impacted by the quality of the read alignment performed. In this manuscript, all indel detection analyses were performed after running BWA-MEM for alignment.

A significant advantage to benchmarking new algorithms is the recent publication and sharing of “reference” genomes derived from long read sequencing. These build on consensus datasets produced by the Genome in a Bottle consortium, which continues to expand its own reference collection in this direction. Notably, basic comparison of these “gold standard” callsets (NA12878 and Syndip) reveals major differences in the number and size distribution of variants too large to be explained by the diversity of individual human genomes. While the Genome in a Bottle initiative and others have made commendable efforts towards creating a complete genome that can be used for benchmarking and tool development, challenges still remain in areas of the genome that are difficult to sequence and characterize; Sanger sequencing can help disambiguate some challenging regions of the genome, but others will require long-read sequencing and other new technologies in order to be completely characterized. Syndip’s use of long-read sequencing and multiple orthogonal variant callers provides for a greater number of variants that span a wider range of sizes, thus offering more comprehensive benchmarking opportunities.

Scotch’s base-by-base procedure is less dependent on indel size than more coarse-grained approaches. For a caller that identifies variants by reconstructing regions of the genome through local assembly, the difference between a 40 bp and a 400 bp indel is significant. But while the complete presentation in sequencing data of these variants may differ, their breakpoints are described by similar combinations of soft-clipped reads and changes in sequencing depth and quality. This relative conformity is the basis of Scotch’s ability to detect indels of drastically different sizes. While trained primarily on NA12878 with a truth set produced by the Genome in a Bottle consortium, we added many large simulated indels to Scotch’s training data to increase its sensitivity to large variants. Though trained only on indels of up to 500 bp, Scotch identifies variants in Syndip of up to several thousand base pairs in length.

We developed a meta caller (Metal) that delivers superior performance overall by integrating five variant callers. Collating the variants reported by these callers in its “smart intersection,” Metal maximizes the number of true positive calls retained while filtering out erroneous calls resulting from sequencing errors. The high number of callers and their high initial sensitivity—as well as the loose comparison requirements—produces a master callset with high recall, including capture of many large variants, and high precision. Across all indel breakpoints in Syndip, Metal’s recall by count is only slightly behind that of Pindel-L and Scotch, which has the highest recall by count of any tool, while greatly improving on Pindel-L and Scotch’s precision.

Sanger sequencing of variants called by Scotch missing from the NA12878 truth set reveals that some “false positives” are *bona fide* indels. Most of these are variants in homopolymer runs. While the reliability of Sanger sequencing itself declines in such regions, the prevalence of variants increases. We view improving sensitivity to these and other broader categories of variants as an imperative. We note that the incompleteness of benchmark datasets, in addition to presenting a challenge to machine-learning based approaches that may learn to miss the same indels, warps metrics derived from benchmarking. Considering the new true positives predicted by Sanger sequencing improves estimates of Scotch’s precision, and may lower estimates of other callers’ recall.

While gold standard datasets provide critical insight into the performance of variant callers, their potential for incompleteness means they should not be relied on exclusively. This is especially true in light of efforts to expand the capabilities of variant callers into broader categories of genetic variants—and those that lie in more challenging genomic regions—where current gold standard datasets are particularly likely to be incomplete. Over-reliance on benchmarking metrics may hinder the development of new tools by incorrectly penalizing improved callers with low precision, and rewarding those that maintain the “status quo” of primarily identifying indels that are already confidently detected. In addition to evaluating callers against benchmark dataset, we encourage evaluation by Sanger sequencing of a sample of calls made outside the truth set for a more full picture of indel callers’ capabilities. While attention to a variety of metrics is important, we urge greater focus on recall and improvement in discovery of a wider range of variants, relative to precision.

## Conclusion

We present Scotch and Metal, tools capable of identifying new true positive insertion and deletion breakpoints, expanding the range of variants that can be detected from next-generation sequencing data. Scotch is intended for maximum recall and identifies more true positive indel breakpoints in Syndip than any other tool examined. Metal integrates the output of Scotch and four other variant callers, capturing many long variants and retaining high sensitivity while improving precision. We also examine gold standard datasets, and show that Sanger sequencing validates some “false positive” variants called by Scotch missing from the NA12878. We hope these tools, aided by insights from benchmark datasets, can continue to advance understanding of human disease and genetic diversity.

## Methods

### Scotch

Scotch is a random forest based indel caller. The pipeline considers each locus individually, computing metrics that describe the sequencing data aligned there and the composition of the reference genome. A random forest model, trained on labeled training data that includes NA12878 variants as well as simulated indels, then classifies each position as a kind of indel breakpoint or non-indel.

#### Input

Scotch accepts a Binary Alignment Mapping (BAM)^21^ file containing next-generation whole-genome sequencing data. Scotch also accepts a FASTA file providing the corresponding reference genome, and BED files delineating the regions of interest. Scotch divides the input by chromosome for parallel processing.

#### Features

Scotch’s calculates 40 features to describe each position, which are later input to a random forest model that classifies each position. Scotch’s features include “primary metrics,” quantities which are extracted directly from sequencing data; “delta features” which track the differences in primary features between neighboring positions; and “genomic features,” which describe the content of the reference genome at a given locus.

#### Primary features

These 12 features are calculated directly from the sequencing data for each locus:

- The number of reads, normalized across the sample (depthNorm)
- The number of reads excluding soft-clipping, normalized across the sample (nReadsNorm)
- Proportion of high quality (base quality ≥ 13) reads to all reads (nHQual)
- Mean base quality (baseQ)
- Mean mapping quality (mapQ)
- Proportion of reads whose CIGAR string indicates an insertion (insRatio)
- Proportion of reads whose CIGAR string indicates soft-clipping (allSCRatio)
- Proportion of reads whose CIGAR string indicates the boundary of a deletion (i.e., the base is deleted but at least one neighboring base is not) (edgeDelRatio)
- Proportion of reads whose CIGAR string indicates the boundary of soft clipping (i.e., the base is soft-clipped but at least one neighboring base is not) (edgeSCRatio)
- Mean base quality of soft-clipped bases (scQual)
- Mean base-pair distance to closest soft-clipped base (scDist)
- *Consistency score* of soft-clipping, if any, defined below (scCons)

A position’s consistency score is a metric we derived that gives the ratio of the number of reads supporting the most common soft-clipped base (i.e., A, T, C, or G), to the number of all soft-clipped reads. Soft-clipping provides important signal of an indel. This score helps a model distinguish indel-related soft-clipping (where all soft-clipped reads should support the same nucleotide) from that caused by low sequencing quality (where different nucleotides will be present).

#### Delta features

These 20 features give the difference, from the locus of interest to each of its neighbors, in each of the primary features listed above, except the soft-clipping consistency score (scCons) and the proportion of high-quality reads (nHQual). These delta features help Scotch, which primarily operates on a base-by-base basis, “see” the context of a given locus to make a more accurate determination of whether it is the site of an indel breakpoint.

#### Genomic features

These 8 features describe the reference genome, providing Scotch with insight into regions where sequencing errors are more common. They include 4 binary features: presence in high-confidence regions, “superdup” regions, RepeatMasker-masked regions^37^, and low-complexity regions. The remaining 4 features are continuous: GC-content within 50bp, GC-content within 1000 bp, mappability, and uniqueness. More information on these features and how they are calculated is available in the Supplementary Note, and on the GitHub pages where they can be downloaded: https://github.com/AshleyLab/scotch-data-grch37 for GRCh37 and https://github.com/AshleyLab/scotch-data-grch38 for GRCh38.

More information on the features and their importance is available in the Supplementary Note (Supplementary Fig. 2, Supplementary Table 19).

#### Output

The features are combined in a TSV that can serve as the input to any number of machine-learning setups. Scotch’s random forest models analyzes the data to classify loci as indel sites or non-indel. Scotch outputs a VCF file that includes all breakpoints discovered.

#### Runtime

Scotch’s runtime is approximately 24 hours, when parallelized by chromosome. Metal’s runtime is approximately 10 minutes, when parallelized by chromosome.

#### Training

We evaluated models’ performance when trained with data from five different sources: simulated variants, Syndip, CHM1, NA12878, and NA12878 with simulated variants. We found optimal results with the last source of training data. We also performed a hyperparameter optimization over random forest hyperparameters including the number of trees (ntree) and the number of predictors that can be considered at each node (mtry), though we found these to be largely insignificant.

### Metal

The meta caller Metal performs a “smart intersection” of calls made by Scotch, DeepVariant, GATK HaplotypeCaller, and Pindel-L. For each of these tools, it decomposes reported indels into a list of breakpoints, each labeled as a deletion start, deletion end, or insertion. Metal reports a breakpoint called by one tool if there is a breakpoint call of the same type by another tool within 3 bp. Metal reports insertions called by Scotch or Pindel-L only if they are within 3 bp of insertion calls by higher-precision DeepVariant, GATK HaplotypeCaller, or VarScan2. Metal outputs a VCF file that includes all breakpoints corroborated in this way.

### F(n) metrics balance recall and precision for clinical variant calling

An F-score considers recall, by base or by count, and precision. An F1-score computes the harmonic mean of recall and precision, giving each equal weight. In an F(N)-score, recall is considered N times more times more important than precision (see Methods).

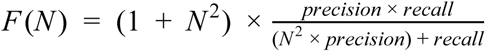

## Declarations

### Ethics approval and consent to participate

This study conforms to the Declaration of Helsinki and was performed as part of the Undiagnosed Diseases Network, which operates under a common IRB protocol.

### Consent for publication

Patients involved in this study provided informed consent.

### Availability of data and material

The codebase is publicly available on GitHub at https://github.com/AshleyLab/scotch, and the genomic features used by the machine learning model are available at https://github.com/AshleyLab/scotch-data-grch37 for GRCh37 and https://github.com/AshleyLab/scotch-data-grch38 for GRCh38.

### Authors& contributions

These authors contributed equally: Charles Curnin, Rachel L. Goldfeder. C.C. implemented the pipeline, analyzed its performance, and drafted the manuscript. R.L.G. developed the approach including the initial pipeline and feature selection. S.M., D.W., M.T.W., and E.A.A., provided statistical and computational support and supervision. R.L.G., C.C., D.W., M.T.W., and E.A.A. designed the experiments and approach. R.L.G., M.T.W., and E.A.A. secured funding. All authors provided critical review and input into the final content of the manuscript.

## Affiliations

*Division of Cardiovascular Medicine, Stanford School of Medicine, Stanford, CA, USA*

Charles Curnin, Shruti Marwaha, Daryl Waggott, Matthew T. Wheeler & Euan A. Ashley

*Jackson Laboratory for Genomic Medicine, Farmington, CT, USA*

Rachel L. Goldfeder

*Stanford Center for Undiagnosed Diseases, Stanford, CA, USA*

Charles Curnin, Shruti Marwaha, Daryl Waggott, Matthew T. Wheeler & Euan A. Ashley

*Department of Genetics, Stanford School of Medicine, Stanford, CA, USA*

Euan A. Ashley

## Consortia

Undiagnosed Diseases Network

Maria T. Acosta, David R. Adams, Pankaj Agrawal, Mercedes E. Alejandro, Patrick Allard, Euan A. Ashley, Mahshid S. Azamian, Carlos A. Bacino, Guney Bademci, Eva Baker, Ashok Balasubramanyam, Dustin Baldridge, Deborah Barbouth, Gabriel F. Batzli, Alan H. Beggs, Hugo J. Bellen, Jonathan A. Bernstein, Gerard T. Berry, Anna Bican, David P. Bick, Camille L. Birch, Stephanie Bivona, Carsten Bonnenmann, Devon Bonner, Braden E. Boone, Bret L. Bostwick, Lauren C. Briere, Elly Brokamp, Donna M. Brown, Matthew Brush, Elizabeth A. Burke, Lindsay C. Burrage, Manish J. Butte, Olveen Carrasquillo, Ta Chen Peter Chang, Hsiao-Tuan Chao, Gary D. Clark, Terra R. Coakley, Laurel A. Cobban, Joy D. Cogan, F. Sessions Cole, Heather A. Colley, Cynthia M. Cooper, Heidi Cope, William J. Craigen, Precilla D&Souza, Surendra Dasari, Mariska Davids, Jean M. Davidson, Jyoti G. Dayal, Esteban C. Dell&Angelica, Shweta U. Dhar, Naghmeh Dorrani, Daniel C. Dorset, Emilie D. Douine, David D. Draper, Annika M. Dries, Laura Duncan, David J. Eckstein, Lisa T. Emrick, Christine M. Eng, Gregory M. Enns, Cecilia Esteves, Tyra Estwick, Liliana Fernandez, Carlos Ferreira, Elizabeth L. Fieg, Paul G. Fisher, Brent L. Fogel, Irman Forghani, Noah D. Friedman, William A. Gahl, Rena A. Godfrey, Alica M. Goldman, David B. Goldstein, Jean-Philippe F. Gourdine, Alana Grajewski, Catherine A. Groden, Andrea L. Gropman, Melissa Haendel, Rizwan Hamid, Neil A. Hanchard, Nichole Hayes, Frances High, Ingrid A. Holm, Jason Hom, Alden Huang, Yong Huang, Rosario Isasi, Fariha Jamal, Yong-hui Jiang, Jean M. Johnston, Angela L. Jones, Lefkothea Karaviti, Emily G. Kelley, Dana Kiley, David M. Koeller, Isaac S. Kohane, Jennefer N. Kohler, Deborah Krakow, Donna M. Krasnewich, Susan Korrick, Mary Koziura, Joel B. Krier, Jennifer E. Kyle, Seema R. Lalani, Byron Lam, Brendan C. Lanpher, Ian R. Lanza, C. Christopher Lau, Jozef Lazar, Kimberly LeBlanc, Brendan H. Lee, Hane Lee, Roy Levitt, Shawn E. Levy, Richard A. Lewis, Sharyn A. Lincoln, Pengfei Liu, Xue Zhong Liu, Sandra K. Loo, Joseph Loscalzo, Richard L. Maas, Ellen F. Macnamara, Calum A. MacRae, Valerie V. Maduro, Marta M. Majcherska, May Christine V. Malicdan, Laura A. Mamounas, Teri A. Manolio, Thomas C. Markello, Ronit Marom, Martin G. Martin, Julian A. Martínez-Agosto, Shruti Marwaha, Thomas May, Jacob McCauley, Allyn McConkie-Rosell, Colleen E. McCormack, Alexa T. McCray, Jason D. Merker, Thomas O. Metz, Matthew Might, Eva Morava-Kozicz, Paolo M. Moretti, Marie Morimoto, John J. Mulvihill, David R. Murdock, Avi Nath, Stan F. Nelson, J. Scott Newberry, John H. Newman, Sarah K. Nicholas, Donna Novacic, Devin Oglesbee, James P. Orengo, Stephen Pak, J. Carl Pallais, Christina GS. Palmer, Jeanette C. Papp, Neil H. Parker, John A. Phillips III, Jennifer E. Posey, John H. Postlethwait, Lorraine Potocki, Barbara N. Pusey, Genecee Renteria, Chloe M. Reuter, Lynette Rives, Amy K. Robertson, Lance H. Rodan, Jill A. Rosenfeld, Robb K. Rowley, Ralph Sacco, Jacinda B. Sampson, Susan L. Samson, Mario Saporta, Judy Schaechter, Timothy Schedl, Kelly Schoch, Daryl A. Scott, Lisa Shakachite, Prashant Sharma, Vandana Shashi, Kathleen Shields, Jimann Shin, Rebecca Signer, Catherine H. Sillari, Edwin K. Silverman, Janet S. Sinsheimer, Kathy Sisco, Kevin S. Smith, Lilianna Solnica-Krezel, Rebecca C. Spillmann, Joan M. Stoler, Nicholas Stong, Jennifer A. Sullivan, David A. Sweetser, Cecelia P. Tamburro, Queenie K.-G. Tan, Mustafa Tekin, Fred Telischi, Willa Thorson, Cynthia J. Tifft, Camilo Toro, Alyssa A. Tran, Tiina K. Urv, Tiphanie P. Vogel, Daryl M. Waggott, Colleen E. Wahl, Nicole M. Walley, Chris A. Walsh, Melissa Walker, Jennifer Wambach, Jijun Wan, Lee-kai Wang, Michael F. Wangler, Patricia A. Ward, Katrina M. Waters, Bobbie-Jo M. Webb-Robertson, Daniel Wegner, Monte Westerfield, Matthew T. Wheeler, Anastasia L. Wise, Lynne A. Wolfe, Jeremy D. Woods, Elizabeth A. Worthey, Shinya Yamamoto, John Yang, Amanda J. Yoon, Guoyun Yu, Diane B. Zastrow, Chunli Zhao, Stephan Zuchner.

