## Supplemental Text for "Machine learning-based detection of insertions and deletions in the human genome"

### Supplemental Appendix

### Full Benchmarking Results

*Results on Syndip*
*Supplementary Table 1: Performance, Syndip, Deletions*

| Tool | Recall by Count | Recall by Base | Precision |
|---|---|---|---|
| DeepVariant | 86.6% | 32.8% | 92.9% |
| GATK HC | 86.1% | 24.0% | 89.3% |
| VarScan2 | 72.7% | 13.9% | 97.0% |
| Pindel | 88.5% | 79.5% | 27.0% |
| Pindel-L | 88.5% | 79.5% | 27.0% |
| Scotch | 89.0% | 56.8% | 76.3% |
| Metal | 90.0% | 60.1% | 84.4% |

*Supplementary Table 2: Performance, Syndip, Insertions*

| Tool | Recall by Count | Recall by Base | Precision |
|---|---|---|---|
| DeepVariant | 85.2% | 31.3% | 93.1% |
| GATK HC | 85.6% | 26.1% | 91.3% |
| VarScan2 | 69.7% | 14.3% | 96.9% |
| Pindel | 12.1% | 2.7% | 14.1% |
| Pindel-L | 89.9% | 40.3% | 2.6% |
| Scotch | 91.7% | 63.9% | 21.0% |
| Metal | 85.4% | 34.6% | 90.5% |

*Supplementary Table 3: Performance, Syndip, All Breakpoints*

| Tool | Recall by Count | Recall by Base | Precision |
|---|---|---|---|
| DeepVariant | 87.1% | 36.0% | 94.6% |
| GATK HC | 87.0% | 25.7% | 91.4% |
| VarScan2 | 73.4% | 14.7% | 97.6% |
| Pindel | 66.3% | 59.8% | 26.9% |
| Pindel-L | 90.6% | 70.5% | 6.2% |
| Scotch | 93.4% | 67.9% | 34.7% |
| Metal | 90.2% | 56.0% | 88.7% |

*Results on Simulated Data*

*Supplementary Table 4: Performance, Simulated Variants, Deletions*

| Tool | Recall by Count | Recall by Base | Precision |
|---|---|---|---|
| DeepVariant | 10.0% | 3.3% | 21.1% |
| GATK HC | 12.1% | 1.2% | 22.2% |
| VarScan2 | 1.3% | 0.0% | 32.1% |
| Pindel | 97.2% | 95.8% | 4.4% |
| Pindel-L | 97.2% | 95.8% | 4.4% |
| Scotch | 97.9% | 98.9% | 38.2% |
| Metal | 90.1% | 88.7% | 32.5% |

*Supplementary Table 5: Performance, Simulated Variants, Insertions*

| Tool | Recall by Count | Recall by Base | Precision |
|---|---|---|---|
| DeepVariant | 4.0% | 2.4% | 20.0% |
| GATK HC | 24.2% | 17.7% | 37.4% |
| VarScan2 | 0.3% | 0.0% | 17.7% |
| Pindel | 1.3% | 0.3% | 0.3% |
| Pindel-L | 41.8% | 40.5% | 0.2% |
| Scotch | 98.1% | 97.4% | 10.7% |
| Metal | 27.0% | 19.2% | 20.1% |

*Supplementary Table 6: Performance, Simulated Variants, All Breakpoints*

| Tool | Recall by Count | Recall by Base | Precision |
|---|---|---|---|
| DeepVariant | 9.0% | 4.7% | 21.3% |
| GATK HC | 22.2% | 14.3% | 42.5% |
| VarScan2 | 1.0% | 0.0% | 29.6% |
| Pindel | 66.2% | 57.5% | 4.1% |
| Pindel-L | 79.4% | 74.0% | 1.0% |
| Scotch | 99.3% | 99.2% | 18.2% |
| Metal | 69.3% | 60.5% | 30.0% |

*Results on NA12878*

*Supplementary Table 7: Performance, NA12878, Deletions*

| Tool | Recall by Count | Recall by Base | Precision |
|---|---|---|---|

| | | | |
|---|---|---|---|
| DeepVariant | 97.8% | 98.3% | 88.8% |
| GATK HC | 97.2% | 97.6% | 88.8% |
| VarScan2 | 84.1% | 85.3% | 98.8% |
| Pindel | 98.7% | 98.1% | 17.1% |
| Pindel-L | 98.7% | 98.1% | 17.1% |
| Scotch | 98.4% | 98.6% | 75.6% |
| Metal | 98.5% | 98.9% | 64.7% |

*Supplementary Table 8: Performance, NA12878, Insertions*

| Tool | Recall by Count | Recall by Base | Precision |
|---|---|---|---|
| DeepVariant | 93.9% | 96.5% | 92.2% |
| GATK HC | 96.4% | 97.1% | 92.5% |
| VarScan2 | 81.4% | 74.8% | 99.4% |
| Pindel | 7.5% | 6.2% | 5.5% |
| Pindel-L | 99.0% | 98.9% | 2.1% |
| Scotch | 97.3% | 97.5% | 24.3% |
| Metal | 94.1% | 98.3% | 71.3% |

*Supplementary Table 9: Performance, NA12878, All Breakpoints*

| Tool | Recall by Count | Recall by Base | Precision |
|---|---|---|---|
| DeepVariant | 96.7% | 98.1% | 90.5% |
| GATK HC | 97.5% | 97.5% | 90.1% |
| VarScan2 | 84.4% | 81.9% | 99.0% |
| Pindel | 78.0% | 74.4% | 19.1% |
| Pindel-L | 99.0% | 98.6% | 5.9% |
| Scotch | 99.2% | 98.6% | 42.5% |
| Metal | 98.7% | 98.8% | 67.4% |

*F Metrics, All Breakpoints*

*Supplementary Table 10: F1 Metrics*

| Tool | Recall by count | Recall by base |
|---|---|---|
| DeepVariant | 90.7% | 52.2% |
| GATK HC | 89.1% | 40.1% |
| VarScan2 | 83.8% | 25.5% |
| Pindel | 38.2% | 37.1% |

| | | |
|---|---|---|
| Pindel-L | 11.7% | 11.5% |
| Scotch | 50.6% | 45.9% |
| Metal | 89.4% | 68.7% |

*Supplementary Table 11: F3 Metrics*

| Tool | Recall by count | Recall by base |
|---|---|---|
| DeepVariant | 87.8% | 38.4% |
| GATK HC | 87.4% | 27.7% |
| VarScan2 | 75.3% | 16.1% |
| Pindel | 57.8% | 53.3% |
| Pindel-L | 38.5% | 34.7% |
| Scotch | 79.9% | 62.0% |
| Metal | 90.1% | 58.2% |

*Supplementary Table 12: F5 Metrics*

| Tool | Recall by count | Recall by base |
|---|---|---|
| DeepVariant | 87.3% | 36.9% |
| GATK HC | 87.2% | 26.4% |
| VarScan2 | 74.1% | 15.2% |
| Pindel | 62.7% | 57.1% |
| Pindel-L | 59.6% | 50.5% |
| Scotch | 87.7% | 65.5% |
| Metal | 90.2% | 56.8% |

*Syndip Recall by Size*

*Supplementary Table 13: Recall by Indel Size, Syndip, Deletions*

| Center Indel Size (bp) | 2.5 | 4.5 | 5.5 | 8 | 15 | 35 | 75 | 300 | 750 | 5500 |
|---|---|---|---|---|---|---|---|---|---|---|
| (N) | 3861 | 838 | 192 | 563 | 557 | 409 | 118 | 95 | 12 | 25 |
| Metal | 0.91 | 1.00 | 0.98 | 0.96 | 0.95 | 0.87 | 0.53 | 0.31 | 0.42 | 0.56 |
| Scotch | 0.91 | 0.99 | 0.97 | 0.95 | 0.95 | 0.74 | 0.37 | 0.20 | 0.42 | 0.56 |
| GATK HC | 0.89 | 0.93 | 0.93 | 0.90 | 0.89 | 0.79 | 0.34 | 0.19 | 0.00 | 0.04 |
| DeepVariant | 0.91 | 0.98 | 0.95 | 0.95 | 0.92 | 0.78 | 0.45 | 0.20 | 0.00 | 0.20 |
| VarScan2 | 0.86 | 0.86 | 0.81 | 0.69 | 0.55 | 0.28 | 0.08 | 0.02 | 0.00 | 0.04 |
| Pindel | 0.89 | 0.93 | 0.95 | 0.91 | 0.93 | 0.91 | 0.65 | 0.51 | 0.67 | 0.80 |
| Pindel-L | 0.89 | 0.93 | 0.95 | 0.91 | 0.93 | 0.91 | 0.65 | 0.51 | 0.67 | 0.80 |

*Supplementary Table 14: Recall by Indel Size, Syndip, Insertions*

| Center Indel Size (bp) | 2.5 | 4.5 | 5.5 | 8 | 15 | 35 | 75 | 300 | 750 | 5500 |
|---|---|---|---|---|---|---|---|---|---|---|
| (N) | 1925 | 390 | 117 | 262 | 261 | 180 | 48 | 72 | 13 | 6 |
| Metal | 0.96 | 0.94 | 0.93 | 0.92 | 0.87 | 0.78 | 0.42 | 0.19 | 0.15 | 0.33 |
| Scotch | 0.96 | 0.96 | 0.98 | 0.95 | 0.93 | 0.73 | 0.48 | 0.50 | 0.38 | 0.83 |
| GATK HC | 0.94 | 0.92 | 0.90 | 0.86 | 0.82 | 0.74 | 0.35 | 0.14 | 0.08 | 0.00 |
| DeepVariant | 0.95 | 0.92 | 0.91 | 0.87 | 0.85 | 0.68 | 0.23 | 0.14 | 0.15 | 0.33 |
| VarScan2 | 0.90 | 0.75 | 0.73 | 0.65 | 0.48 | 0.18 | 0.00 | 0.07 | 0.00 | 0.00 |
| Pindel | 0.19 | 0.08 | 0.03 | 0.08 | 0.07 | 0.04 | 0.00 | 0.04 | 0.00 | 0.00 |
| Pindel-L | 0.94 | 0.93 | 0.94 | 0.91 | 0.89 | 0.86 | 0.67 | 0.35 | 0.46 | 0.00 |

*Supplementary Table 15: Recall by Indel Size, Syndip, All Breakpoints*

| Center Indel Size (bp) | 2.5 | 4.5 | 5.5 | 8 | 15 | 35 | 75 | 300 | 750 | 5500 |
|---|---|---|---|---|---|---|---|---|---|---|
| (N) | 5786 | 1228 | 309 | 825 | 818 | 589 | 166 | 167 | 25 | 31 |
| Metal | 0.93 | 0.98 | 0.96 | 0.95 | 0.94 | 0.86 | 0.51 | 0.29 | 0.36 | 0.55 |
| Scotch | 0.95 | 0.99 | 0.98 | 0.98 | 0.98 | 0.86 | 0.60 | 0.47 | 0.60 | 0.68 |
| GATK HC | 0.91 | 0.93 | 0.92 | 0.91 | 0.90 | 0.80 | 0.34 | 0.18 | 0.08 | 0.03 |
| DeepVariant | 0.93 | 0.96 | 0.94 | 0.94 | 0.92 | 0.77 | 0.43 | 0.22 | 0.16 | 0.26 |
| VarScan2 | 0.88 | 0.83 | 0.78 | 0.69 | 0.53 | 0.27 | 0.08 | 0.05 | 0.00 | 0.03 |
| Pindel | 0.68 | 0.69 | 0.63 | 0.68 | 0.67 | 0.67 | 0.51 | 0.32 | 0.40 | 0.68 |
| Pindel-L | 0.92 | 0.95 | 0.96 | 0.93 | 0.93 | 0.92 | 0.68 | 0.47 | 0.56 | 0.71 |

*Simulated Variants Recall by Size*

*Supplementary Table 16: Recall by Indel Size, Simulated Variants, Deletions*

| Center Indel Size (bp) | 7.5 | 15 | 35 | 75 | 300 | 750 | 1750 | 6250 |
|---|---|---|---|---|---|---|---|---|
| (N) | 14 | 24 | 66 | 112 | 590 | 859 | 144 | 170 |
| Metal | 1.00 | 1.00 | 1.00 | 0.99 | 0.93 | 0.91 | 0.60 | 0.93 |
| Scotch | 1.00 | 1.00 | 0.91 | 0.84 | 0.99 | 0.99 | 1.00 | 0.99 |
| GATK HC | 1.00 | 1.00 | 1.00 | 1.00 | 0.02 | 0.01 | 0.01 | 0.00 |
| DeepVariant | 1.00 | 1.00 | 1.00 | 0.98 | 0.10 | 0.03 | 0.03 | 0.02 |
| VarScan2 | 0.57 | 0.58 | 0.06 | 0.00 | 0.00 | 0.00 | 0.00 | 0.00 |
| Pindel | 1.00 | 1.00 | 1.00 | 1.00 | 1.00 | 1.00 | 0.65 | 0.99 |

| | | | | | | | | |
|---|---|---|---|---|---|---|---|---|
| Pindel-L | 1.00 | 1.00 | 1.00 | 1.00 | 1.00 | 1.00 | 0.65 | 0.99 |

*Supplementary Table 17: Recall by Indel Size, Simulated Variants, Insertions*

| Center Indel Size (bp) | 7.5 | 15 | 35 | 75 | 300 | 750 | 1750 | 6250 |
|---|---|---|---|---|---|---|---|---|
| (N) | 7 | 13 | 32 | 60 | 287 | 408 | 41 | 144 |
| Metal | 1.00 | 0.92 | 0.91 | 0.55 | 0.22 | 0.22 | 0.20 | 0.18 |
| Scotch | 1.00 | 1.00 | 0.97 | 0.88 | 0.99 | 0.99 | 0.98 | 0.97 |
| GATK HC | 1.00 | 0.92 | 1.00 | 0.47 | 0.18 | 0.19 | 0.20 | 0.17 |
| DeepVariant | 1.00 | 0.92 | 0.94 | 0.37 | 0.05 | 0.05 | 0.02 | 0.01 |
| VarScan2 | 0.29 | 0.08 | 0.00 | 0.00 | 0.00 | 0.00 | 0.00 | 0.00 |
| Pindel | 0.00 | 0.15 | 0.16 | 0.02 | 0.01 | 0.01 | 0.00 | 0.00 |
| Pindel-L | 1.00 | 1.00 | 1.00 | 0.67 | 0.32 | 0.38 | 0.32 | 0.44 |

*Supplementary Table 18: Recall by Indel Size, Simulated Variants, All Breakpoints*

| Center Indel Size (bp) | 7.5 | 15 | 35 | 75 | 300 | 750 | 1750 | 6250 |
|---|---|---|---|---|---|---|---|---|
| (N) | 21 | 37 | 98 | 172 | 877 | 1267 | 185 | 314 |
| Metal | 1.00 | 0.97 | 0.97 | 0.85 | 0.70 | 0.69 | 0.53 | 0.59 |
| Scotch | 1.00 | 1.00 | 0.98 | 0.99 | 0.99 | 0.99 | 1.00 | 0.99 |
| GATK HC | 1.00 | 0.97 | 1.00 | 0.81 | 0.14 | 0.13 | 0.12 | 0.14 |
| DeepVariant | 1.00 | 0.97 | 0.98 | 0.77 | 0.10 | 0.05 | 0.05 | 0.04 |
| VarScan2 | 0.48 | 0.41 | 0.04 | 0.00 | 0.00 | 0.00 | 0.00 | 0.00 |
| Pindel | 0.67 | 0.70 | 0.74 | 0.70 | 0.69 | 0.68 | 0.51 | 0.54 |
| Pindel-L | 1.00 | 1.00 | 1.00 | 0.90 | 0.78 | 0.80 | 0.60 | 0.74 |

### Recall by Indel Size in Syndip

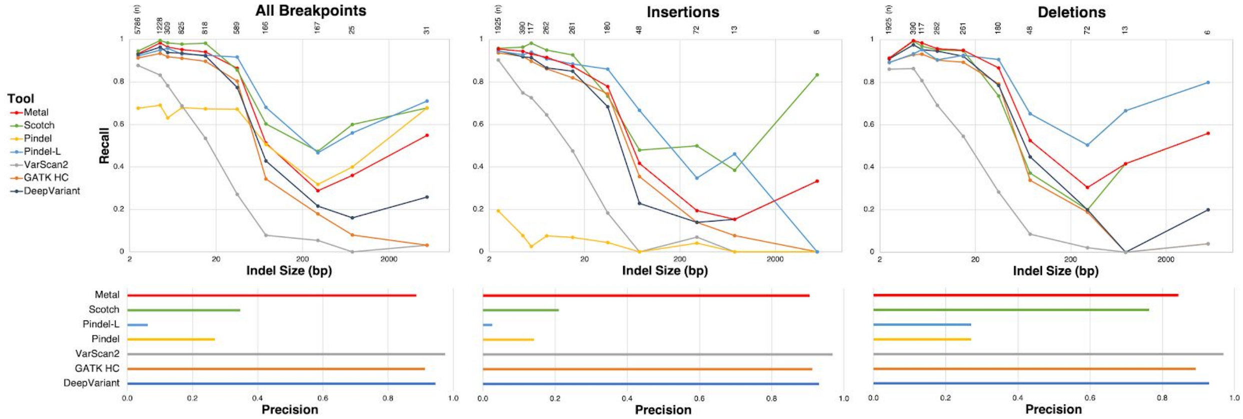


*Supplementary Fig. 1:* Recall by indel size and precision by caller in Syndip.
Scotch and Pindel-L are capable of identifying large variants other tools miss, though at the expense of lower precision. Scotch is more successful than many other indel callers in identifying larger insertions. Metal, a meta caller comprising the other indel callers navigates a compromise in performance, falling just short of Scotch's recall at some size categories but gaining much in precision.

### Features

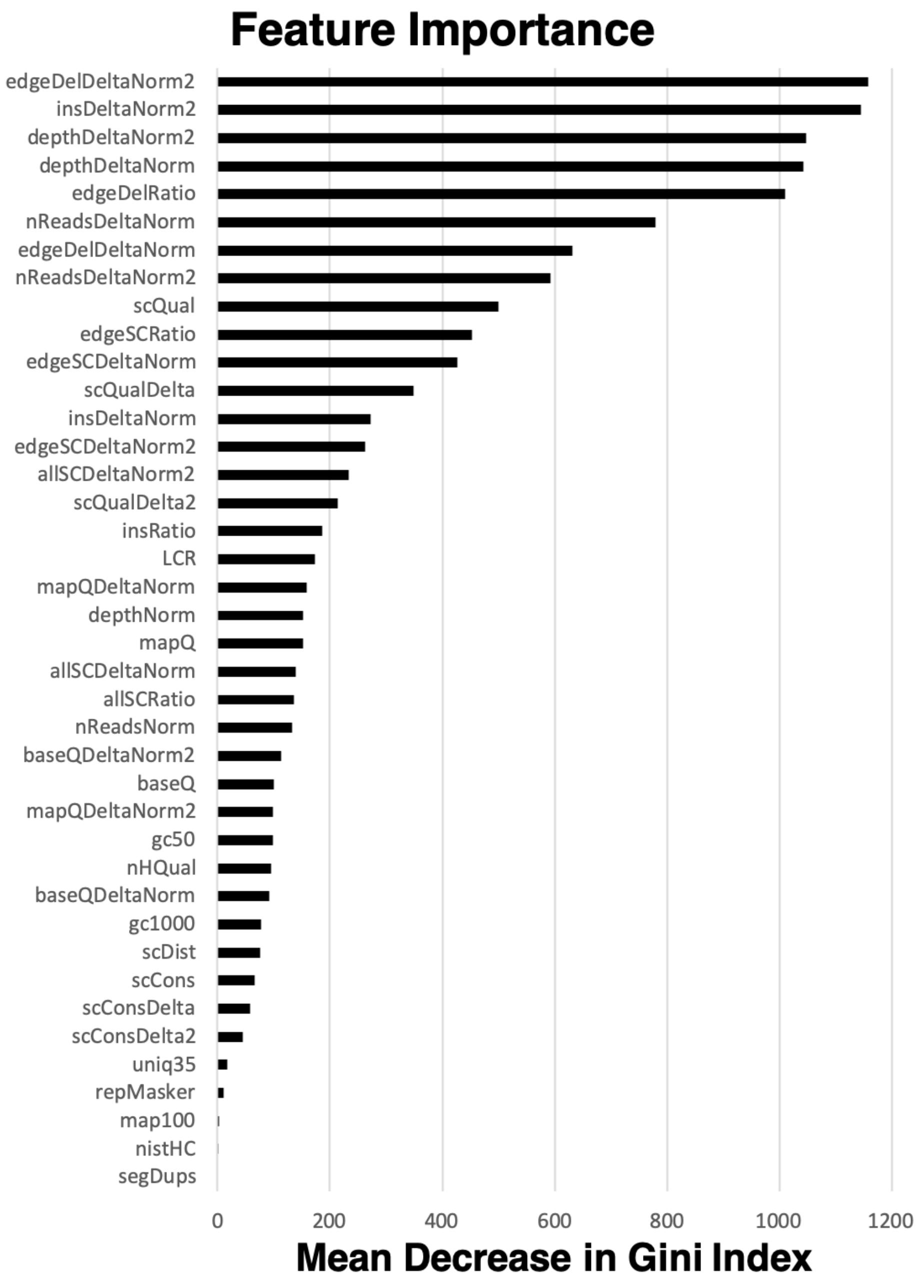


*Supplementary Fig. 2*: Feature Importance
We provide the mean decrease in Gini index for each feature used in our random forest model. Many important features come from analyzing reads' CIGAR string, which describes their alignment to the reference. While Scotch first calculates features for a specific position, it also calculates "delta" metrics that describe the difference in these features between the given position and adjacent ones. These provide particularly valuable signal to the model.

*Supplementary Table 19: Description of Features*

| Name | Mean decrease in Gini index | Description | Calculated from |
|---|---|---|---|
| edgeDelDeltaNorm2 | 1157.432308 | change in edgeDelRatio | primary features |
| insDeltaNorm2 | 1145.035001 | change in insRatio | primary features |
| depthDeltaNorm2 | 1046.858365 | change in depthNorm | primary features |
| depthDeltaNorm | 1042.792823 | change in depthNorm | primary features |
| edgeDelRatio | 1010.55461 | proportion of reads with "D" boundary at this position | CIGAR string |
| nReadsDeltaNorm | 779.6781336 | change in nReadsNorm | primary features |
| edgeDelDeltaNorm | 631.815455 | change in edgeDelRatio | primary features |
| nReadsDeltaNorm2 | 593.2460816 | change in nReadsNorm | primary features |
| scQual | 499.9683733 | mean base quality of SC bases | custom metric |
| edgeSCRatio | 452.3754421 | proportion of reads where this position is first or last SC base | CIGAR string |
| edgeSCDeltaNorm | 425.9807946 | change in edgeSCRatio | primary features |
| scQualDelta | 348.6274247 | change in scQual | primary features |
| insDeltaNorm | 272.7460816 | change in insRatio | primary features |
| edgeSCDeltaNorm2 | 263.233626 | change in edgeSCRatio | primary features |
| allSCDeltaNorm2 | 232.8962805 | change in allSCRatio | primary features |
| scQualDelta2 | 214.4870579 | change in scQual | primary features |
| insRatio | 185.5387626 | proportion of reads with "I" at this position | CIGAR string |
| LCR | 172.9696488 | whether in low-complexity region acc. to GIAB | reference genome |
| mapQDeltaNorm | 158.8558588 | change in mapQ | primary features |
| depthNorm | 151.9115883 | coverage | pileups |

| | | | |
|---|---|---|---|
| mapQ | 151.746711 | mapping quality | quality score |
| allSCDeltaNorm | 138.5197639 | change in allSCRatio | primary features |
| allSCRatio | 135.8175727 | proportion of reads with SC | CIGAR string |
| nReadsNorm | 133.167538 | coverage (reads with no SC) | pileups |
| baseQDeltaNorm2 | 113.0688671 | change in baseQ | primary features |
| baseQ | 100.3728414 | base quality | quality score |
| mapQDeltaNorm2 | 98.5384296 | change in mapQ | primary features |
| gc50 | 98.99349585 | GC content within 50 bp window | reference genome |
| nHQual | 95.4076284 | proportion of reads with alignment quality ≥ 13 | pileups |
| baseQDeltaNorm | 91.27668783 | change in baseQ | primary features |
| gc1000 | 77.24163302 | GC content within 1000 bp window | reference genome |
| scDist | 75.41347764 | base pair distance to closest soft clipped base | custom metric |
| scCons | 65.67716663 | defined above | custom metric |
| scConsDelta | 57.61233851 | change in scCons | primary features |
| scConsDelta2 | 44.33997025 | change in scCons | primary features |
| uniq35 | 17.49447824 | UCSC Genome Browser: wgEncodeDukeMapabilityUniqueness35bp | reference genome |
| repMasker | 10.41891019 | whether masked by RepeatMasker | reference genome |
| map100 | 2.993679202 | UCSC Genome Browser: wgEncodeCrgMapabilityAlign100mer | reference genome |
| nistHC | 0.1102200571 | whether in NIST high-confidence regions | reference genome |
| segDups | 0 | whether in segmental duplication region acc. to GIAB | reference genome |

*Genomic features*

Eight features Scotch analyzes describe the reference genome. Using the abbreviated names in Supplementary Table 2, they are nistHC, repMasker, segDups, LCR, gc50, gc1000, map100 and uniq35. The first four features are binary and discrete; the last four vary continuously.

- nistHC indicates whether the position lies in high-confidence regions identified by NIST Genome in a Bottle (GiaB)[1]; the underlying data is available at ftp://ftp-trace.ncbi.nlm.nih.gov/giab/ftp/release/NA12878_HG001/NISTv3.3.2/GRCh37/HG001_GRCh37_GIAB_highconf_CG-IllFB-IllGATKHC-Ion-10X-SOLID_CHROM1-X_v.3.3.2_highconf_nosomaticdel.bed
- repMasker indicates whether the position lies in RepeatMasker[2]-masked regions; the underlying data is available in the UCSC Genome Browser[3] "rmsk" table
- segDups indicates whether the position lies in GiaB segmental duplications; the underlying data is available at ftp://ftp-trace.ncbi.nlm.nih.gov/giab/ftp/data/NA12878/analysis/NIST_union_callsets_06172013/superdupsmerged_all_sort.bed.gz
- LCR indicates whether the position lies in known low-complexity regions[4]; the underlying data is available at https://figshare.com/articles/Low_complexity_regions_in_hs37d5/969685
- gc50 gives the GC content of the position within a 50-bp window; the data is calculated with the "bedtools nuc"[5] command
- gc1000 gives the GC content of the position within a 1000-bp window; the data is calculated with the "bedtools nuc" command
- map100 gives a mappability score for the position; the underlying data is available through the UCSC Genome Browser "wgEncodeCrgMapabilityAlign100mer" table[6]
- uniq35 gives a uniqueness score for the position; the underlying data is available through the UCSC Genome Browser "wgEncodeDukeMapabilityUniqueness35bp" table

**Metrics**

Indel calling is a problem of highly imbalanced classes. For every position that should be identified as an indel breakpoint, there are on the order of ten thousand that should not be identified as indel breakpoints. A classified could thus achieve 99.99% accuracy by simply "predicting" that every position was not an indel. Such a classifier would miss every mutation and have little utility, however.

For this reason, we express callers' performance through the metrics of recall (sensitivity) and precision (positive predictive value). The hypothetical classifier would score a zero on both. Recall is the ratio of true positives to true positives and false negatives; precision is the ratio of true positives to true positives and false positives. These quantities capture, roughly, how likely a real variant is to be called, and how likely a called variant is to be real, respectively.

The traditional definition of recall (or "recall by count") is calculated with respect to the number of variants that are true positives or false negatives. A classifier that identifies a 1 bp deletion but fails to identify a 3 bp deletion would score a recall of 50%. Because of the preponderance of small variants, an indel pipeline can achieve high recall just by identifying small variants, systematically missing larger ones. To examine callers' performance in identifying indels across the size spectrum, we introduce the metric of recall by base, which is calculated with respect to the number of bases belonging to the variants identified. A classifier that identifies a 1 bp deletion but fails to identify a 3 bp deletion would score a recall by base of only 25%.

**Training**

Our training configuration consists of four parameters: 1) a training dataset, 2) a corresponding "truth" providing the location of all known indels, 3) a set of features calculated to describe the dataset, and 4) certain fixed hyperparameters for training a model. We examined four datasets, with three different versions of truth files for each; five different features sets; and eight different hyperparameter configurations.

*Datasets*
Training a machine-learning model to recognize indels requires a genome in which the indels are already known. Datasets like this can come from two sources: 1) published "gold standard" genomes, like NA12878, Syndip and CHM1; or 2) data simulation, in which a computational tool can be directed precisely to "spike-in" designated mutations at given positions. BAMSurgeon, used in the DREAM Somatic Mutation Calling Challenge, is one such tool. Starting with sequencing data for NA12878, we excluded all portions of the genome at which indels had been identified by GIAB, thus obtaining a kind of "clean slate." We defined 1,488 indels—three for each size [5, 500) bp—at random positions and random zygosities. We trained models relying on NA12878, Syndip, CHM1, simulated variants, and NA12878 with simulated variants. We found the best results from the last.

*Truths*
Each of the four datasets of known indels has an associated truth set, which should characterize the indels present. Even with our simulated data, however, we noted many of the variants appeared to be located at positions slightly different from what was recorded in the VCF. The soft-clipping or drop in coverage associated with a deletion, for example, might begin one base pair after the position recorded in the truth. This discrepancy can have a serious effect on lowering the quality of Scotch's model. Because we use the truth to label individual positions as indel breakpoints, if a variant actually covers positions different from what is recorded in the truth set, this item in the training data will encourage Scotch to identify a normal site as a breakpoint, and a breakpoint as a normal site.

To address this systematically, we defined a heuristic for each breakpoint: for the left breakpoints of deletions, the change in coverage from the previous position to the current position; for the right breakpoints of deletions, the change in coverage from the current position to the next position; for insertion sites, the number of reads that begin to be soft-clipped at the position. For each breakpoint recorded in the truth, we re-assigned the indel position' to the locus within 3 bp with the highest heuristic.

*Features*
We also explored five different sets of features to identify which set of predictors yielded the best performance. Scotch performed best when trained with a moderate number of features tested

(40). With additional predictors that characterized the existing features of a given position relative to more distal loci, performance declined. This suggest that the features selected can accurately identify breakpoints by characterizing just the given locus and its nearest neighbors.

*Hyperparameters*

Random forest models are collections of decisions trees. Each tree is in turn a collection of nodes, which represent conditions that involve predictors. Depending on which conditions are satisfied, each tree will “vote” for a given class, like “deletion start breakpoint” or “insertion.” The model’s final output is the class for which the most trees vote.

A random forest has two hyperparameters: the number of trees in the forest (ntree), and the number of predictors that can be considered at each node (mtry). We considered four possible values for ntree (350, 450, 550, 650), and two for mtry (7, 8), resulting in eight possible hyperparameter configurations. We found these to be largely insignificant, resulting in little charge in the performance of the resultant model.

*Reporting alleles*

A disadvantage of Scotch’s base-by-base approach is that except in the case of 1-bp deletions, Scotch does not report the nucleotide sequence of the alternate allele. Instead, it provides “<INS>,” “<DEL_L>,” or “<DEL_R>”: insertion site, start of a deletion, or end of a deletion. To recover the full variant allele, this output can be directed to specialized tools that perform read assembly. For insertions large relative to read length, reconstructing the full inserted sequence involves assembling not just local reads, but also “orphan” reads that were not mapped to the indel site because they have so few reference bases. The format of Scotch’s reports are in general similar to versions of Pindel, which for many variants report “<INS>” or “<DEL>.”

There may be deletions for which Scotch can identify only one of the two breakpoints. In this case, a <DEL_L> or <DEL_R> record may appear in the VCF without an obvious mate. Such calls may be false positives; but they may also be real capture of an indel for which one edge is particularly difficult to identify. In keeping with our goal of increasing sensitivity for large indel calling, we do not filter out such calls. While having both breakpoints of a deletion is preferable to having just one, we believe that having even one is preferable to having neither. Even without an alternate allele sequence or a mate breakpoint, knowledge that an indel of a given type exists at specific coordinates is valuable information.

**Metal**

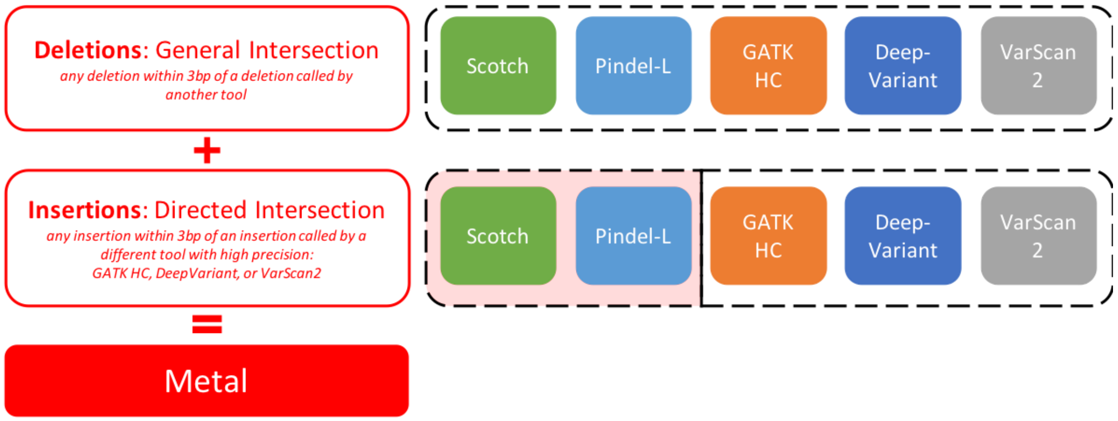


*Supplementary Fig. 3: Metal is a meta-analytic indel caller*
Metal will report a call produced by a tool if it has a corresponding call within 3 bp identified by another tool. To counter the low insertion-specific precision of Scotch and Pindel-L, we require that insertions called by these tools have correlates in higher-precision DeepVariant, GATK HaplotypeCaller, or VarScan2.

### Sanger Sequencing

Of the 18 verified calls, 15 lay in homopolymer stretches. While such regions are known to be hotspots of indels and other mutations, traditional homopolymer indel calling suffers from low sensitivity, which means homopolymer indels may be especially likely to be missing from benchmark datasets[7,8]. Scotch's ability to identify such variants is noteworthy. At the same time, in homopolymer regions both Sanger and next-generation sequencing are less reliable, meaning that even Sanger sequencing may serve as less of a validating "gold standard"[9] and illustrating the overall unsolved challenge of this variant class.

Of the 18 verified calls, 17 were deletions, whereas only 12 were predicted to be deletions. This mirrors the results derived from benchmarking, where Scotch's deletion-specific precision is higher than its insertion-specific precision. Scotch had misclassified 5 of the validated deletions as insertions, which suggests that while Scotch is able to identify the presence of indels in homopolymer regions, it does not perform as well at identifying their type.

Of the 18 validated calls, 2 had correlates in Syndip. (Of the 34 invalid calls, another 3 had correlates in Syndip.) One insertion breakpoint and one deletion breakpoint called in NA12878—which were absent from the NA12878 truth set but supported by Sanger sequencing—were within 3 bp of indel breakpoints in Syndip. This may suggest that Syndip, relative to NA12878, includes some common indel polymorphisms missing from the NA12878 truth set, but miniscule sample size prevents a more confident assertion.

*Supplementary Table 20: Sanger Sequencing Results*

| Position | Predicted Type | Has Syndip Correlate? | Validation Result | Validation Type | In Homop.? |
|---|---|---|---|---|---|
| 17682853 | del_L | no | no | | yes |
| 17866545 | del_R | no | yes | del | yes |
| 18597740 | del_L | no | no | | yes |

| | | | | | |
|---|---|---|---|---|---|
| 19685871 | ins | no | yes | del | no |
| 20055244 | ins | yes | yes | ins | yes |
| 21170227 | del_L | no | yes | del | yes |
| 21173593 | del_R | no | yes | del | yes |
| 25198493 | ins | no | no | | yes |
| 26164161 | ins | no | no | | no |
| 26441040 | ins | no | no | | yes |
| 26896621 | ins | no | yes | del | yes |
| 27038613 | ins | no | no | | no |
| 28773188 | del_L | no | yes | del | yes |
| 29338461 | ins | no | no | | no |
| 29378555 | del_R | no | no | | yes |
| 31815360 | ins | no | yes | del | yes |
| 31962107 | ins | yes | yes | del | yes |
| 32614047 | ins | no | no | | no |
| 32813267 | ins | no | no | | no |
| 33133243 | ins | no | no | | yes |
| 33586415 | ins | no | no | | no |
| 35043111 | ins | no | yes | del | no |
| 36133049 | ins | no | no | | no |
| 36249339 | del_R | no | yes | del | yes |
| 36479840 | ins | no | no | | no |
| 36577730 | del_R | no | yes | del | no |
| 36691447 | del_L | no | no | | no |
| 39400226 | del_L | no | no | | yes |
| 39752475 | ins | no | no | | no |
| 39785958 | del_L | no | yes | del | yes |
| 39972811 | del_L | no | no | | yes |
| 40338627 | del_L | no | yes | del | yes |
| 40452802 | del_R | no | yes | del | yes |
| 40516340 | del_R | no | yes | del | yes |
| 40644082 | del_R | no | yes | del | yes |
| 40807998 | ins | no | no | | yes |
| 41191301 | del_L | yes | no | | yes |

| | | | | | |
|---|---|---|---|---|---|
| 41264434 | del_R | no | no | | no |
| 41382911 | ins | no | no | | yes |
| 41387823 | ins | no | no | | yes |
| 44197707 | ins | no | no | | yes |
| 45149106 | del_R | no | no | | yes |
| 46126224 | del_L | yes | no | | yes |
| 46545641 | del_L | no | no | | no |
| 47342388 | ins | no | no | | no |
| 47994307 | del_R | yes | no | | yes |
| 48450264 | del_L | no | yes | del | yes |
| 48689187 | ins | no | no | | no |
| 49413932 | del_L | no | no | | yes |
| 49665151 | ins | no | no | | yes |
| 49748828 | ins | no | no | | no |
| 50356319 | ins | no | no | | no |

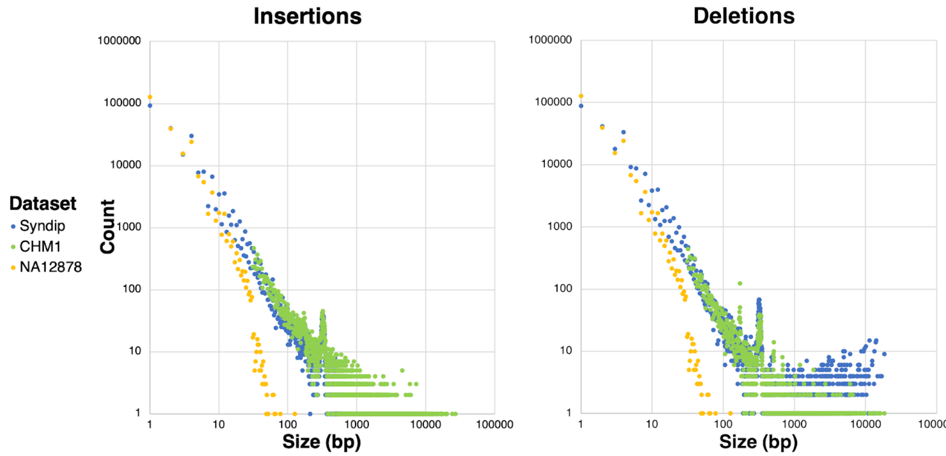


*Supplemental Fig. 4:* Log-scale distribution of indels by size by dataset.
Despite both deriving from human genomes, NA12878 contains fewer indels than Syndip, and smaller ones. We also examine here CHM1[10], a haploid complete hydatidiform mole for which there exists a comprehensive list of indels larger than 30 bp detected by SMRT sequencing. Ostensibly, Syndip, which consists of two cell lines (CHM1 and CHM13), should contain more indels than CHM1 alone. But even within certain size ranges, CHM1 has more variants. While both datasets have a peak in the number of indels around 320 bp (representing SINEs, short interspersed nuclear elements), CHM1 alone has an additional peak around 170 bp[11]. These variants are either false positives in CHM1, or false negatives in Syndip.

**Overview of select cases considered from Undiagnosed Diseases Network**

*Supplementary Table 21: Overview of cases*

| AGE | SEX | Human Phenotype Ontology terms |
|---|---|---|
| 12 | M | HP:0001824; HP:0007338; HP: 0011477; HP: 0025402; HP: 0002650; HP: 0001260; HP: 0100543; HP: 0010862; HP: 0007018; HP: 0001337; HP: 0002015; HP: 0002487; HP: 0007302; HP: 0100034; HP: 0100035; HP: 0002072; HP: 0001618; HP: 0002019 |
| 43 | M | HP:0001824; HP: 0007994; HP: 0002321; HP: 000164; HP: 0001962; HP: 0003202; HP: 0003325; HP: 0003552; HP: 0003710; HP: 0003712; HP: 0007185; HP: 0003401; HP: 0002018; HP: 0003236 |
| 61 | M | HP:0003394,HP:0003403,HP:0003552,HP:0100295,HP:0003548,HP:0000939,HP:0000704,HP:0003165,HP:0003691 |
| 4 | F | HP: 0003508; HP: 0000252; HP: 0001508; HP: 0000324; HP: 0000463; HP: 0000582; HP: 0004453; HP: 0000407; HP: 0000960; HP: 0007616; HP: 0002099; HP: 0001252; HP: 0001276; HP: 0002194; HP: 0001269; HP: 0012469; HP: 0100021; HP: 0011471; HP: 0006266; HP: 0010881 |
| 23 | F | HPO IDs not available |
| 9 | M | HPO IDs not available |
| 31 | F | HP:0003323,HP:0003805,HP:0003458,HP:0003715,HP:0009130,HP:0010628,HP:0003236,HP:0003691,HP:0002111 |
| 18 | F | HP:0001348,HP:0001765,HP:0003401,HP:0007002,HP:0008075,HP:0001884,HP:0007178 |
| 13 | F | HP:0001644 |
| 2 | F | HPO IDs not available |
| 1 | M | HP:0000252, HP:0100704, HP:0001601, HP:0003808, HP:0001263, HP:0001257 |
| 22 | M | HP:0001263,HP:0000252,HP:0002558,HP:0000486,HP:0002942 |
| 6 | M | HPO IDs not available |
| 5 | F | HP:0000577,HP:0000522,HP:0001600,HP:0002500,HP:0001344,HP:0008897,HP:0000574,HP:0011231,HP:0000767 |
| 12 | F | HPO IDs not available |
| 10 | M | HPO IDs not available |
| 5 | M | HP:0001263,HP:0001250,HP:00055484,HP:0001252,HP:0006817,HP:0007370 |
| 73 | M | HPO IDs not available |
| 20 | M | HPO IDs not available |
| 43 | F | HP:0003326, HP:0002829, HP:0000821, HP:0000794, HP:0003493, HP:0025289, HP:0100769, HP:0008180, HP:0100602, HP:0008071 |
| 13 | M | HPO IDs not available |

| | | |
|---|---|---|
| 11 | F | HPO IDs not available |
| 30 | M | HP:0000639, HP:0001962, HP:0012664, HP:0001252, HP:0001324; HP:0003394, HP:0008947, HP0000112, HP:0009830, HP:0002625, HP:0100658, HP:0000093, HP:0003236, HP:0045045 |
| 1 | F | HPO IDs not available |
| 16 | M | HPO IDs not available |
| 37 | M | HPO IDs not available |
